# Atypical Pupil-Linked Arousal Induced by Low-Risk Probabilistic Choices, and Intolerance of Uncertainty in Adults with ASD

**DOI:** 10.1101/2024.04.11.589067

**Authors:** Kristina I. Pultsina, Tatiana A. Stroganova, Galina L. Kozunova, Andrey O. Prokofyev, Aleksandra S. Miasnikova, Anna M. Rytikova, Boris V. Chernyshev

## Abstract

Adults with autism spectrum disorder (ASD) report stress when acting in a familiar probabilistic environment, but the underlying mechanisms are unclear. Their decision-making may be affected by the uncertainty aversion implicated in ASD, and associated with increased autonomic arousal. Previous studies have shown that in neurotypical (NT) people, decisions with predictably better outcomes are less stressful and elicit smaller pupil-linked arousal than those involving random “trial-and-error” searches or self-imposed risk of exploration. Here, in a sample of 46 high-functioning ASD and NT participants, we explored pupil-linked arousal and behavioral performance in a probabilistic reward learning task with a stable advantage of one choice option over the other. Using mixed-effects model analysis, we contrasted pupil dilation response (PDR) between a preferred frequently rewarded exploitative decision and its explorative alternatives. We observed that subjects with ASD learned the advantageous probabilistic choices at the same rate over time and preferred them to the same degree as NT participants both in terms of choice ratio and decision speed. Despite similar reward prediction abilities, outcome predictability modulated decision-related PDR in ASD in the opposite direction than in NT individuals. Moreover, relatively enhanced PDR elicited by exploitative low-risk decisions predicted a greater degree of self-reported intolerance of uncertainty in everyday life. Our results suggest that in a non-volatile probabilistic environment, objectively good predictive abilities in people with ASD are coupled with elevated physiological stress and subjective uncertainty regarding the decisions with the best possible but still uncertain outcome that contributes to their intolerance of uncertainty.

## Main Text

Autism spectrum disorder (ASD) is a neurodevelopmental condition characterized by difficulties with social communication, repetitive behaviors, and limited interests (*Diagnostic and statistical manual of mental disorders : DSM-5™*, 2013). Even those adults with ASD who have normal intellectual abilities have difficulty making everyday decisions in both social and non-social contexts (Luke et al., 2012). Individuals with ASD are known to experience stress and anxiety when they are uncertain about something (Kanner, 1968; Robic et al., 2015), and this intolerance of uncertainty (Hwang et al., 2020) can hinder their ability to pursue their goals by flexibly adapting behavior to statistical patterns in the world.

However, research on the behavior of people with ASD in structured probabilistic environments has yielded mixed results. In multiple choice and binary probabilistic reward tasks, they perform no differently than neurotypical (NT) subjects and learn to favor the option with more frequent rewards at the same rate as NT subjects (reviewed by (Zeif & Yechiam, 2020), but see (South et al., 2014; Vella et al., 2018; Zhang et al., 2015) for results on superior or inferior performance in ASD). Importantly, all of these studies found no group differences in probabilistic reward learning between autistic and non-autistic adult participants when the experimental paradigm did not impose elevated demands on memory, or when feedback was unambiguous and only one explicit rule was required to be followed.

In a recent review on decision-making in autism, van der Plas, Mason, and Happe (van der Plas et al., 2023) highlighted that despite typical behavior in simple probabilistic decision-making tasks, subjective assessment of outcomes in adults with autism (i.e., confidence in their preferred choices) is at odds with their objectively high-performance accuracy. For example, participants with ASD, compared with neurotypical individuals, were less aware of their objective accuracy when assessing their confidence verbally, but despite reduced awareness, they still successfully adjusted their behavior to account for errors (Carpenter et al., 2019; Doenyas et al., 2019; Nicholson et al., 2019). This suggests that rather than having an atypical behavior, adult subjects with autism are less confident in their decisions. Indeed, confidence is defined as a form of metacognition associated with some other subjective human qualities such as introspection and self-awareness, and the latter qualities are often impaired in autism (reviewed by (Hatfield et al., 2019; Huggins et al., 2020)).

The dissociation between behavior and its internal evaluation is a complex phenomenon, not least because it raises the question of why people with autism lack confidence to the extent that they forego the benefits of their successful probabilistic learning and avoid making decisions in probabilistic situations (Luke et al., 2012). In the neuroscience literature, confidence in one’s decision is referred to as an inverse (reciprocal) measure of subjective uncertainty (Meyniel et al., 2015). Subjective uncertainty (uncertainty in decision outcomes) is associated with increased decision stress and is a powerful factor that increases peripheral autonomic arousal, which in turn can be tracked by non-luminance-related changes in pupil size (Critchley, 2005; de Berker et al., 2016).

Here, we used a pupil dilation response (PDR) as a proxy measure of the autonomic arousal generated by decision-making in ASD in a simple two-alternative reward learning task, in which choice-reward contingency was probabilistic.

Decision-making in probabilistic tasks relies on prior experience – participants tend to choose options associated with a higher probability of positive outcomes in the past. In a predictive coding framework (e.g., (Smith et al., 2021)), they form an experience-based prediction model that allows them to maximize the average reward. Experimental manipulations of external factors that decrease confidence in the predicted outcome potentiate decision-related pupil dilation. In particular, pupil size has been found to respond to sudden changes in feedback structure that require updates of the prediction model (e.g., (Kreis et al., 2023; Pajkossy et al., 2023)), to a conflicting choice between two options with equally high values in the past (Van Slooten et al., 2018), or to a conflict between the current decision and its expected adverse consequences (Kozunova et al., 2022). The degree of pupil dilation has been hypothesized to be positively scaled to decision uncertainty (Kreis et al., 2023; Lavin et al., 2013; Satterthwaite et al., 2007). In previous studies with NT participants, the two intertwined aspects of such uncertainty have been assessed at different time intervals of a slowly evolving pupil dilation response (PDR). PDR scaling with decision uncertainty occurs in the time window between explicit choice and external feedback, thereby reflecting an internal state that is unrelated to the evaluation of external information about the outcome of the current choice (de Berker et al., 2016; Poe et al., 2020; Urai et al., 2017). On the other hand, subjective uncertainty is also encoded in PDR driven by the feedback signal itself (Kreis et al., 2023; Pajkossy et al., 2023). Feedback-induced PDR is thought to signal a discrepancy between predicted and actual decision outcomes. Subjective uncertainty about one’s predictive model leads to a large feedback-related PDR, regardless of the feedback valence (unsigned prediction error, or surprise), thereby increasing the amount of new information that has to be assimilated to update the model.

Decision-making in probabilistic reward learning tasks relies on several strategies that maintain the balance between exploration and exploitation and deal with different types of uncertainty (Dayan, 2012). Exploitative decisions pursue a known reward, rely on the prior experience of the best choice (prediction model of choice value), and are relatively safe and effortless. Yet, uncertainty in exploitation-based decisions is inherent in probabilistic relationships between choices and rewards and is called expected uncertainty (Yu & Dayan, 2005). Exploration, on the other hand, involves two different strategies that are applied depending on the prior knowledge of choice-reward conditions. One is random exploration, where an unknown or changing reward probability induces exploration “by chance” to obtain a prediction model for navigating through uncertainty and to guide behavior toward greater reward. Because this risk is unknown and has to be learned through experience, there is estimation uncertainty or ambiguity in every choice. A popular way to induce random exploration is task rule reversals, when actions or choices that used to be rewarded become unproductive or less productive, thus causing “unexpected uncertainty”. Environments associated with excessive unexpected uncertainty are highly stressful since they lack stable relationships and pose substantial potential problems for safe exploitation (Dayan, 2012).

Another form of exploration is directed exploration, in which the probabilities of reward are known and the selection of a less-known or less-chosen option is encouraged by an informational bonus at the expense of missing a reward (Kozunova et al., 2022; Wilson et al., 2014). Therefore, for directed exploration, one’s risk estimation is high (Dayan, 2012) and, as compared with exploitative strategy, increases the subjective uncertainty of the decision-maker.

Previous studies of phasic pupillary response in autism or in people with autistic traits have focused on reversal learning tasks, in which unexpected uncertainty was driven by changes in the rules defining the choice-outcome contingency (Kreis et al., 2023; Lawson et al., 2017). For example, in a study by Lawson and colleagues (Lawson et al., 2017), participants were presented with two auditory cues that were probabilistically associated with different visual images (e.g., 80% and 20%) and asked to recognize the next image as quickly as possible. This probability changed over time, formally defining several task periods with low and high environmental volatility. In this task, individuals with ASD showed atypically diminished discrimination during volatile versus stable periods of the tasks, both in terms of their choice reaction time (RT) and post-outcome pupil dilation. The authors interpreted blunted discriminative responses as indications that ASD subjects experienced a mild surprise due to overestimated environmental volatility independent of the true predictability of events. However, because of frequent transitions from stable to unstable task periods, the reversal learning task caused anxious participants with ASD to be subjectively uncertain about any choice they made and to view all options as risky (Shi et al., 2022). Therefore, the atypical pattern of PDR exhibited by subjects with ASD in the reversal learning paradigm, though informative, does not answer the question of why people with ASD lack confidence in tasks in which the choice-outcome contingency is stable, albeit probabilistic.

The current study aimed to address this gap by examining the autonomic concomitants of exploitative and explorative choices in ASD in a task with stable choice-outcome probabilities. Because the external feedback favored the same choice throughout each block, this study may clarify whether the previously reported non-discriminative pupil response in ASD reflects genuinely increased subjective uncertainty in the prediction of outcomes or stressfulness of the reversal learning task by itself.

Recently, using such a task, we found a strong link between choice type and pupil-related phasic arousal in a large sample of NT participants (Kozunova et al., 2022). We used a two-alternative forced-choice task, in which reward probability was substantially different between the two alternatives (the objectively advantageous choice had a much higher reward probability than the disadvantageous choice: 70% vs. 30%). The participants learned choice-reward contingency from monetary feedback, with only one explicit instruction requiring the participants to maximize their monetary gain. When probabilities are unknown to the subject, uncertainty is high for both alternatives, hence increasing pupil-related post-decision arousal and choice reaction time. After the prediction model is formed, the objectively advantageous alternative becomes the preferred choice with faster decision time and smaller pupil dilation response. Nevertheless, participants still make rare disadvantageous choices, probably in a purposeful search for information, at the cost of not using the option that is currently judged to be better. In favor of this assumption was the fact that risky exploratory versus safe exploitative decisions were associated with simultaneous increases in post-decision pupil dilation, increased decision-related brain activation, and slower decision times (Chernyshev et al., 2023; Kozunova et al., 2022). Thus, neurotypical subjects can optimally scale their subjective uncertainty, i.e., reduce decision time, brain activity, and autonomic arousal when choosing an option that has been frequently rewarded in the past (i.e., exploitative choice), while maintaining high phasic arousal and prolonged decision time when making a deliberate risky choice that conflicts with the model-based prediction (i.e., a directed exploratory choice). Importantly, this simple task does not leave room for unexpected uncertainty, because safe exploitative, and directed explorative risky actions are under control of the decision maker, who chooses which strategy to follow.

Here, using the same paradigm, we directly tested the idea that intellectually high-functioning adults with ASD will experience atypically high subjective uncertainty in their predictive model under a stable non-volatile environment and this will be reflected in their pupil-linked arousal. Under the assumption of subjectively increased expected uncertainty, we anticipated that ASD individuals would exhibit atypically reduced differences in phasic pupil dilation between model-congruent exploitative and model-incongruent risky explorative choices. In addition, we expected that the objective autonomic measures of stress driven by decision uncertainty in ASD participants would positively correlate with self-reported intolerance of uncertainty they experienced in everyday life.

## Methods

### Participants

Twenty-three neurotypical (NT) volunteers and twenty-three people diagnosed with autism spectrum disorders (ASD), matched for age (neurotypical: mean age 28.17 years, age range 19–42 years; participants with ASD: mean age 29.22 years, age range 20–44 years) and gender (5 men and 18 women in each group), took part in the experiment (Table 1). Eighteen NT subjects were selected from a larger sample of 89 subjects, who participated in our previous pupil study (Kozunova et al., 2022), while additional 5 NT subjects were recruited to match the ASD individuals for age and gender. All participants reported no neurological disorders and had normal or corrected to normal vision. The diagnosis of ASD was established according to the DSM–V criteria of autism spectrum disorder (*Diagnostic and statistical manual of mental disorders : DSM-5™*, 2013) by a board-certified psychiatrist experienced in evaluating ASD and comorbid psychiatric disorders, who is the co-author of this paper (N.A.Y). In addition, the full version (50 items) of the Autism Quotient (AQ) (Baron-Cohen et al., 2001) questionnaire was completed by participants with ASD for the assessment of self-reported autistic traits. The mean AQ total score in this group was 41.20 (AQ range 22–48, SD. 6.70). Two of the 23 participants with a diagnosis of ASD verified by a clinician did not reach the recommended cut-off of 32 cumulative points in the AQ total score, but as they met the other criteria of ASD, they were not excluded.

**Table 1.** Demographics.

|  | NT group | ASD group | *p* |
| --- | --- | --- | --- |
| Number of participants | 23 | 23 | — |
| Male/Female number | 5/18 | 5/18 | ns |
| Age (years) | 28.2 ± 6.0 | 29.2 ± 6.9 | ns |
| Wechsler Performance IQ | 106.8 ± 7.0 | 104.6 ± 8.9 | ns |
| Autism-spectrum quotient | — | 41.2 ± 6.7 | — |

The Russian translation of the Wechsler Performance IQ (PIQ) (WAIS, third edition, UK) was administered to assess intellectual functioning (Table 1). ASD and NT participants did not differ in PIQ (Mann–Whitney U =311, n1=n2= 23, p=0.311 two-tailed). The study was conducted in accordance with the Helsinki Declaration and was approved by the local Ethics Committee of the Moscow State University of Psychology and Education. All participants provided written informed consent.

### Intolerance of Uncertainty (IU)

Intolerance of Uncertainty (IU) refers to the tendency to respond negatively to uncertain or unexpected situations and events in terms of emotions, cognition, and behavior (Stuart et al., 2020). IU is linked to key ASD features, such as insistence on sameness, inflexibility in routines, and avoidance of changes (Joyce et al., 2017; South & Rodgers, 2017). Participants were administered the New Tolerance-Intolerance of uncertainty questionnaire designed and validated for Russian-speaking respondents (Kornilova, 2010). The Intolerance of Uncertainty subscale which we used in our study includes 13 statements (e.g. “The best managers have instructions so precise that subordinates have nothing to worry about” or “It is better to stick to the chosen way of doing things than to change it, because this can lead to confusion”) evaluating the trait-like predisposition to perceive and interpret uncertain situations as triggering negative expectations, excessive worrying, and avoidance of such situations. Based on their responses, participants were assigned individual scores using a seven-point scale ranging from 1 (not at all typical of me) to 7 (entirely typical of me). Altogether, the maximal score could reach 13*7=81 points (highly negative attitude towards uncertainty) whereas the smallest rating is equal to 13*1=13 points. Six NT participants did not have Intolerance of Uncertainty scores.

### Experimental design

The task and procedure were the same as in the previous study where they were reported in detail (Kozunova et al., 2022). Briefly, a modified two-alternative probabilistic learning task (Frank et al., 2004; Kozunova et al., 2022) was offered to the participants in the form of a computer game. Experimental presentation and stimulation were controlled by Presentation software (Presentation 14.4, Neurobehavioral Systems, Albany, USA). The experiment contained 6 blocks, 40 trials each. A short rest of 1 min (or longer, if a participant required) separated the blocks. Participants were instructed that there was an underlying contingency rule between the stimulus and the reward, and their task was to maximize the reward they received over time through trial-and-error experience. In every trial, participants chose between the two stimuli presented on the screen simultaneously and received feedback in the form of points she/he got. Within each block, one of the stimuli was associated with a greater probability of better outcomes than the other (gains on 70% of trials in blocks 1–5, and 60% of trials in block 6). In each block, participants encountered a new pair of stimuli and chose repeatedly between them. Probabilities did not change within any blocks of the experiment. Losses and wins were assigned to each stimulus within a new pair in a quasi-random order. We used the following reinforcement schemes with varying magnitude of wins and losses: (I) +20 & 0, (II) 0 & −20, (III) +50 & +20, (IV) −20 & −50, (V) +20 & −20, (VI) −20 & +20. The rationale behind changing reinforcement schemes was to reduce participants’ propensity to get bored because of the repetition of the same task. We counterbalanced the order of the schemes over the blocks using the following sequences: I–II–III–IV–V–VI, V–III–II–IV–I–VI, and V–III–II–I–IV–VI, each allowing keeping the total score of gains and losses at positive rates to avoid frustration and demotivation in participants. For each participant, we assigned the block sequences randomly. At the end of each block, participants were shown their total score accumulated over the course of the block. At the end of the experiment, we converted the accumulated score to rubles at a 1:1 ratio and paid the participants.

### Trial structure

In every trial, the stimulus pair was presented following the fixation cross (which lasted 150 ms) and was rendered in white on a black background; the stimulus pair remained on the screen until a participant pressed a button. After termination of the decision process (indicated by the button press), the stimuli disappeared, and the black screen was empty until a feedback signal occurred with a delay of 1000 ms, and then it remained on the screen for 500 ms. The inter-trial interval varied pseudo-randomly between 700–1200 ms. Each of the two stimuli within a pair was the same Hiragana hieroglyph (visual angle 1.54 × 1.44°) rotated at two different angles and presented to the left and to the right from the fixation cross at 1.5° of visual angle (Fig. 1). Location of the advantageous stimulus on the left or right side of the screen was alternated pseudo-randomly over trials within each block. The feedback images presented on each trial showed points earned on a current trial. We also made sure that different stimulus pairs as well as feedback images for different earning levels had the same size and contrast border length across six blocks. This was important for the evaluation of luminance-independent pupil-related arousal because the pupil reacts not only to luminance but also to spatial features of the retinal image (Young et al., 1993).

**Figure 1.**
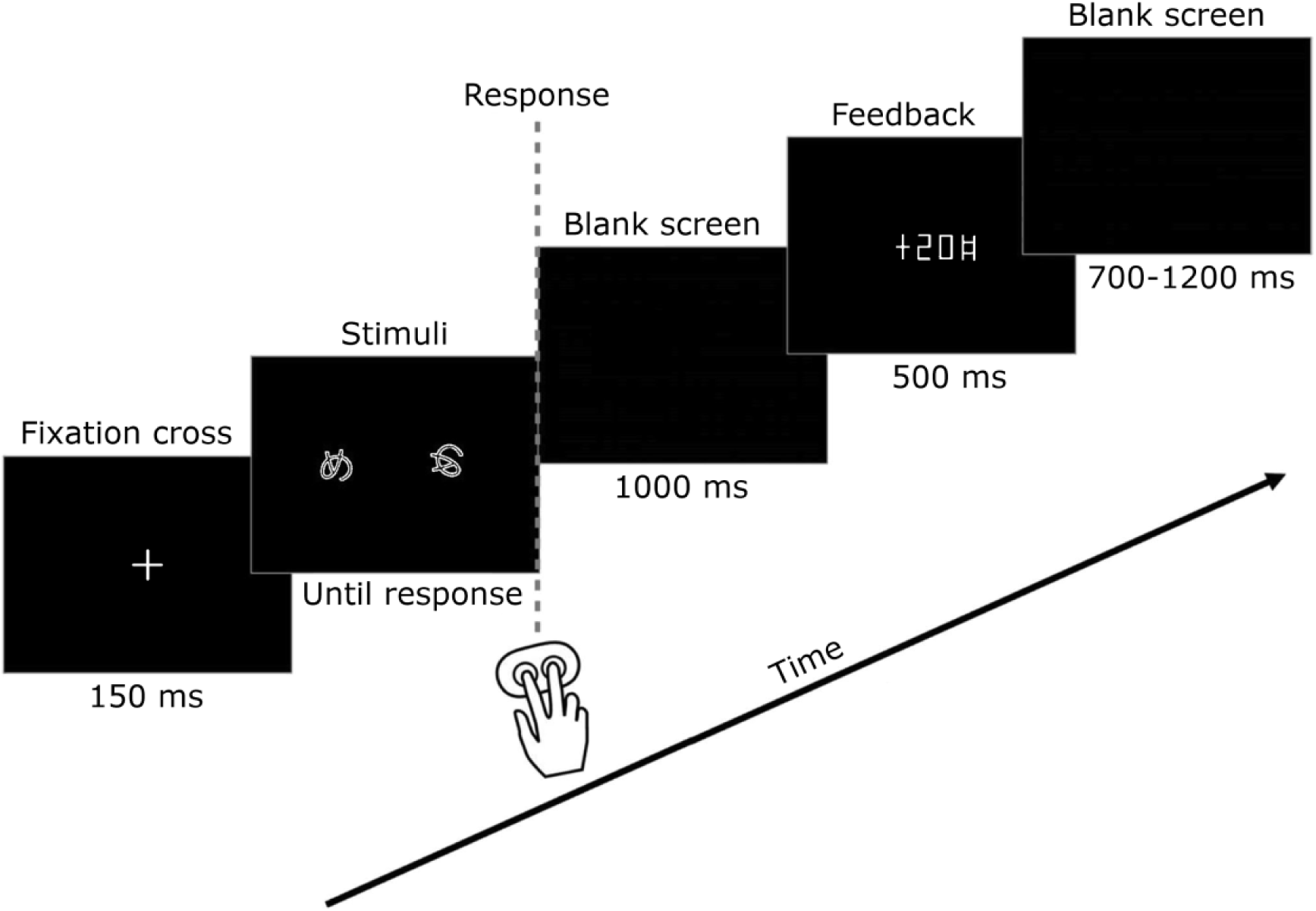
Experimental procedure. Participants learned to select the more profitable of the two images (the advantageous option) solely from probabilistic feedback. After each trial, the number of points earned by the participant was displayed after a 1,000-ms delay after a participant chose one of the options by pressing the left or right button (for the left and right choices respectively). See text for details.

### Procedure

Over the course of the experiment, participants were seated in a quiet and darkened room, in front of a semi-opaque screen, on which the image was projected by Panasonic PT-D7700E-K projector with a resolution of 1280−1024. Participants’ heads were placed on the chinrest to avoid involuntary head movements, and the participants kept their index and middle fingers of the right hand on two buttons of the response pad (Current Designs, Philadelphia, Inc., PA). Eye level and camera position remained constant throughout the recording session for each participant. When choosing the stimuli, participants pressed the correspondent button, i.e., they pressed the left button to choose the stimulus on the left, and they pushed the right button to choose the stimulus on the right.

### Pupillometric data recording

The pupil size of the participant’s dominant eye was measured using an infrared eye tracker EyeLink 1000 Plus (SR Research Ltd., Canada) with a sampling rate of 1,000 Hz and default eye tracker settings. Before each block, participants completed the EyeLink 1000 9-point calibration procedure.

### Task performance and pupillometry

Task performance was assessed as a proportion of objectively advantageous and disadvantageous choices and choice reaction time, which were determined by an iterative algorithm that assigned button responses to stimulus events. For each button response, the RT in milliseconds to the last stimulus pair was calculated. For both behavioral and RT analysis, we excluded trials with extremely low (< 300 ms) and extremely high (> 4,000 ms) RT values resulting in discarding 9.5% of the data. Further, RT values were log-transformed to make the distribution closer to normal.

For offline analyses of pupil data, we developed customized scripts in R Studio (Version 4.2.1 R Core Team, 2022) using the “eyelinker” package (Barthelme, 2019). We identified and removed artifactual data caused by blinks or deficient recording. When doing so, we used the same trial rejection criteria as in the previous research: a whole 5-sec epoch (−2000 to 3000 ms relative to the button press) was removed if pupillometric data were missing or the absolute value of the derivative of the pupillometric data exceeded 10 pixels between two adjacent measurements for longer than 350 ms. If the artefactual pupillometric data segments were shorter than 350 ms (which we considered eyeblinks), we replaced the data using linear interpolation. Overall, this led to discarding pupil data for 22% of epochs. No participants were excluded from the further analysis.

Further, we downsampled pupil data to 20 time points per second by averaging the data within every consecutive 50-ms interval. Then the data were time-locked to the corresponding button press and epoched from 2 seconds before to 3 seconds after the button press. The epochs in which pupil size fell beyond 3*sigma from the mean pupil sizes in the target time interval of 1000–2200 ms were deleted. This step resulted in additional discarding of 0.1% of the data.

For each subject, we applied z-transformation to pupillary data across “clean” epochs from all six experimental blocks. The mean and standard deviation for pupil size were calculated over all epochs and over all time points. The z-scores for every time point of the epoch reflected the number of standard deviations from the common reference value. As discussed by (Kozunova et al., 2022), z-normalization allows comparisons of pupil size between experimental conditions in the same temporal window or between temporal windows in the same condition, with larger z-score value corresponding to greater pupil size. Unlike conventional baseline normalization, which is blind to the risk of a carryover effect from a preceding trial (see (Attard-Johnson et al., 2019)), z-score time courses preserve information on a relative pupil size at the trial onset thus showing its contribution to the ensuing decision-related pupil response. Of note, z-score normalization eliminates any absolute difference in pupil size between ASD and NT groups for any certain condition/choice type and only allows evaluation of the effect of group membership on *relative* between-condition variation in pupil size.

### Trial selection criteria and factors of interest

To examine the general pattern of pupil size and RT measures for exploitative and directed explorative choices in NT and ASD participants, we employed the same approach as in our previous research (Kozunova et al., 2022). These two types of choices can be distinguished after the formation of a predictive model, which is formed by way of extrapolation from regularities extracted from the preceding trial-and-error experience, and such a model can bias decisions towards objectively advantageous choices. Therefore, for every participant and in each block, we first identified task periods belonging to the “after learning” experimental condition, i.e., after a strong preference for the more rewarded stimulus was formed. The following criteria were applied: 1) within each block, “after learning” periods followed four uninterrupted successive advantageous choices; 2) from this moment and until block termination, the percentage of advantageous choices was greater than 65% (i.e., higher than 50% probability, p<0.05, one-tailed binomial test).

To differentiate between exploitative and directed explorative choices under “after learning” condition, we defined an exploitative choice as an objectively advantageous choice surrounded by advantageous choices (“high-payoff”, or HP), and a directed explorative one as a disadvantageous one (“low-payoff”, or LP) surrounded by advantageous HP choices (Table 2) (factor Choice type in analyses described below). The rationale for placing exploitative choice within a sequence of HP choices is the sensitivity of pupil size not only to the directed explorative disadvantageous choice itself but also to preceding and following advantageous choices (Kozunova et al., 2022).

**Table 2.**
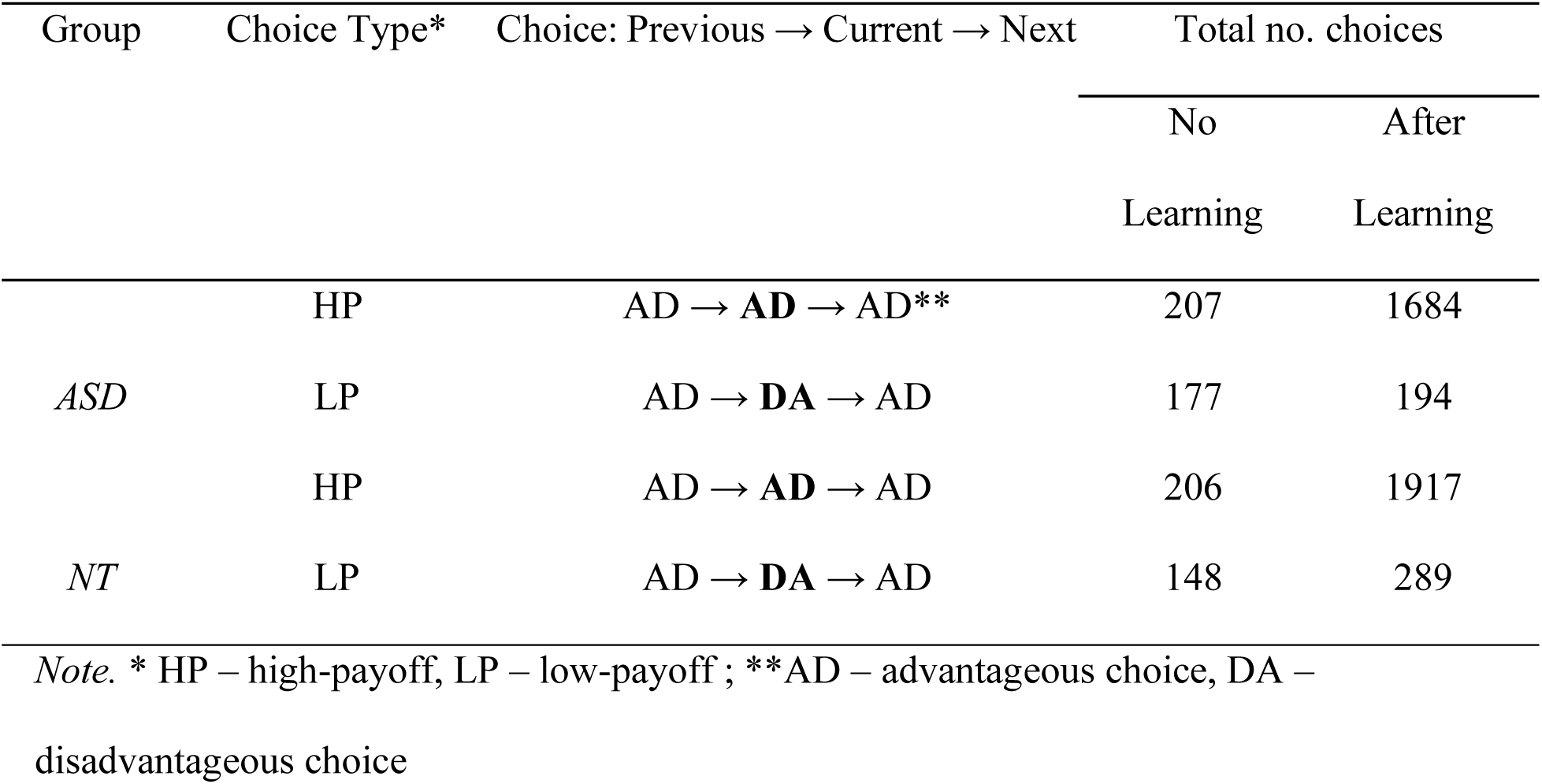
Choice types and the number of epochs sampled for linear mixed-effects model (LMM) analysis.

Those blocks that did not meet performance criteria and in which a participant did not select a preferred alternative by the end of the block (40 trials) were considered as “no learning” condition, where no choice-reward association was ever established. For the sake of uniformity during comparisons, we applied the same technical rules to extract HP and LP trials from “no learning” condition as from “after learning” condition.

To unravel the functional consequences of the predictive model acquisition, we compared HP and LP choices between “after learning” and “no learning” experimental conditions (factor Learning). For both conditions, we applied identical rules for choice type identification as described above, but it is important to keep in mind that HP and LP choices under the “no learning” condition did not represent exploitative and explorative strategies per se, both being samples of “random exploration”. Table 2 shows the number of trials sampled for this analysis per each experimental condition and choice type in the ASD and NT groups separately.

Another factor of interest was the impact of feedback obtained during the previous trial on pupil size measured in the current trial (factor Previous Feedback). Earlier we found that in NT participants, the PDR concomitants of a directed exploration were significantly affected by an outcome of an immediately preceding exploitative choice, which provides a piece of external evidence for the validity of the inner utility model and either supports (negative outcome) a following decision to explore an alternative option or counteracts with it (positive outcome) (Kozunova et al., 2022). Here, we evaluated whether ASD and NT participants had the same short-term consequences of previous feedback for pupil-linked arousal in the current trial.

Similarly, we addressed the feedback presented on a current trial (factor Current Feedback).

### The pupil size waveforms and Time Interval of Interest for pupil size measures

To examine the general temporal dynamics of pupil changes and how it differs between exploitative and directed explorative choices, the respective response-locked pupil traces in each time point were compared using the two-tailed permutation tests with FDR correction for multiple comparisons (Benjamini & Hochberg, 1995). For the ASD and NT groups separately, we applied the R *lmp*() function, which fits linear models to the single-trial data assigning a Choice Type as a fixed factor and a subject as a random factor, and we used 1000 permutations to obtain p-values. This approach does not depend on any theoretical assumptions about the data, avoids any experimenter bias associated with choosing a period over which to compute summary statistics, and reveals when the pupil trajectories for exploitative and explorative choices begin to diverge over time in NT and ASD groups.

For all PDR analyses (see below), we chose the temporal interval between 1000 and 2200 ms after the button press (which equals to 0–1200 ms after feedback onset). This selection served a dual purpose: firstly, it aligned with the post-feedback interval assessed in previous pupil studies of ASD in volatile environments (Kreis et al., 2023; Lawson et al., 2017), facilitating comparisons with the prior research; secondly, this interval corresponded to the period during which we previously observed the most pronounced choice-specific pupil response in NT subjects (Kozunova et al., 2022). However, we found a similar yet attenuated significant effect of choice type in the preceding 500 ms interval, i.e., much earlier than feedback was produced (Kozunova et al., 2022). Therefore, it is highly likely that choice-specific pupil response in the interval between 0 and 1200 ms post-feedback onset does not solely reflect feedback-related arousal as was suggested earlier (e.g., (Kreis et al., 2023; Lawson et al., 2017)), but also the arousal related to the decision to choose the desired option. This is especially true because the differential task-related pupil response requires hundreds of milliseconds to develop (Bradley et al., 2017; Isabella et al., 2019).

For each participant, experimental condition, and choice type separately, z-scored values of pupil size were averaged across the designated interval, and the mean value was subjected to linear mixed-effects models (LMM) analysis to test out the hypothesis of suboptimal pupil-related arousal in people with ASD, which is less discriminative between safe exploitative and risky explorative decisions.

### Statistical analysis of RT and pupil size using the linear mixed effects model

We conducted statistical tests using linear mixed-effects model (LMM) at a single-trial level; the LLM method is robust to imbalanced designs (Kliegl et al., 2011). LMM assesses systematic changes of PDR and RT across trials representing different choice types and conditions, delineates consistently present group or condition differences (fixed effects), and controls for different sources of variability in the data, e.g., inter-individual variability between participants and/or between blocks (random effects).

Statistical LMM analyses of RT and pupillometric data were performed using R Studio (Version 4.2.1 R Core Team, 2022) and the lme4 package (Bates et al., 2015). We examined the following fixed effects and their interactions: “Choice Type” (2 levels: HP and LP, see above); “Learning” (2 levels: “no learning”, and “after learning”, as described above); “Previous Feedback” (2 levels: gain, and loss – the outcome of the trial that immediately preceded the current choice); “Group” (2 levels: NT and ASD participants). To control for individual and between-block variability we assigned “Subject” (46 levels), and “Block” (6 levels for the number of blocks in the experimental sequence) to random effects.

The following models were run for the main analyses:

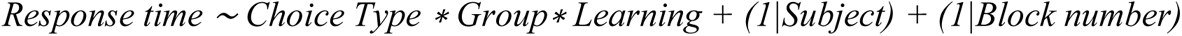

and

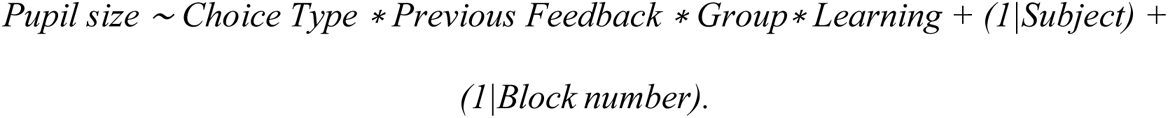

For the elimination of nonsignificant effects (for details see (Kozunova et al., 2022)), we performed a step-down model selection procedure using the “step” function implemented in lmerTest package (Kuznetsova et al., 2017). We applied the Satterthwaite approximation for denominator degrees of freedom and reported results using a type III ANOVA table (package lmerTest, function ANOVA) (Kuznetsova et al., 2017). The estimated marginal means were derived from the results of the Linear Mixed Effects Model (LMM). Subsequently, these means were utilized for both creating visual plots and carrying out Tukey HSD post hoc tests using the emmeans R package (Lenth, 2021).

### Statistical analysis of behavior

To examine the effect of group membership and learning on the propensity of decision-makers to choose one option over the other, we utilized a Generalized Mixed-Effects Logistic Regression Model (GLMM). The model was implemented using the *glmer* function from the R package lme4, with the family parameter set to “binomial” to accommodate for the binary nature of the outcome variable. The dependent variable was expressed as the odds ratio (OR) for advantageous choices (AD), with the fixed factors being group membership (NT and ASD), learning (“no learning”, “after learning”), and we used the following model:

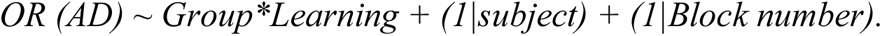

To evaluate the effect of the group membership on the participants’ propensity to switch between two options across the whole experimental session, we calculated the OR for the switch trials (AD to DA or DA to DA), and used the model:

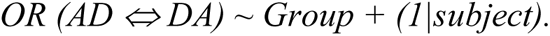

We also assessed the group difference in the effect of feedback valence (gain or loss) in the “after learning” condition on the participants’ tendency to switch from advantageous to disadvantageous option in the next trial:

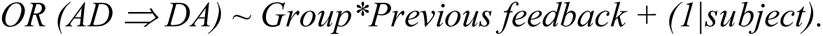

Results were considered significant at p < 0.05, with p-values derived from Wald-Z tests.

### Correlational analysis

To evaluate the relationship between pupil measures and Intolerance of uncertainty scores in our participants, Spearman’s correlation was used. All testing was conducted two-sided with a significance level of 0.05 after Bonferroni correction of p-level for multiple comparisons (number of comparisons=6: 3 Groups – NT, ASD, the whole sample, and 2 choice types – explorative and exploitative).

## Results

### Behavioral performance

The two groups had a similar number of “unsuccessful” blocks in which no learning occurred: 1.8 ± 1.9 blocks out of six in NT and 1.4 ± 1.4 in ASD (ASD vs NT: t (44) = 0.71, p = 0.48). In successful blocks, participants in the NT and ASD groups required a statistically indistinguishable number of attempts out of the 40 trials comprising each block to reach learning criteria: NT: 11.5 ± 12.9, and ASD: 13.9 ± 12.3 (ASD vs NT: t (44) = 0.06, p-value = 0. 949). The NT and ASD participants did not differ in the cumulative payoff over the whole experiment: ASD: 417.1 +/– 353.6 rubles (range –200 – 960), NT: 460.8 +/– 305.9 rubles (range –110 – 1000), ASD vs NT: t (44) = 0.68, p = 0.500.

Table 3 summarizes by-group statistics on the percentage of disadvantageous and advantageous choices split by learning condition (no learning and after learning). The GLMM did not reveal a significant effect of the group on the ratio of AD choices made (OR=0.92, z = – 0.12, p=0.907) or significant interaction between the learning factor and the group (OR=0.95, z = –0.37, p=0.715). Concurrently, the effect of learning on the ratio of AD choices was found to be significant (OR=3.48, z =11.28, p<0.001). Participants in both groups equally displayed a significant bias toward advantageous choices in the “after learning” but not in the “no learning” condition (Table 3).

**Table 3.** Task performance.

| Condition | Group | Proportion by Choice Type |  | Total No. of choices |  |
| --- | --- | --- | --- | --- | --- |
|  |  | *AD* | *DA* | *AD* | *DA* |
| No learning | NT | 52% | 48% | 862 | 786 |
|  | ASD | 51% | 49% | 1012 | 988 |
| After learning | NT | 84% | 16% | 2565 | 495 |
|  | ASD | 85% | 15% | 2141 | 372 |

In addition, we compared groups by choice switching ratio across all choices in six blocks of forty trials each. This was done because some studies reported that adults with ASD tend to exhibit extensive choice switching in the Iowa Gambling Test – a reward learning task with a stable reward structure (Zeif & Yechiam, 2020). Choice switching occurred on average in 31% and 35% of trials in the NT and ASD groups respectively and the ratio did not differ between the groups: GLMM (OR =1, z = 0.1, p = 0.920) indicating that ASD subjects did not have increased propensity for choice switching in our task.

We also checked for putatively increased behavioral sensitivity to the negative feedback in the ASD participants, as a possible source of their atypical behavior in cognitive tasks (see e.g. Broadbent & Stokes, 2013). When the stimulus-reward contingency was learned by NT and ASD participants, respectively, 40% and 43% of all transitions from advantageous to disadvantageous choices were committed after losses. The GLMM has not revealed a significant effect of the previous feedback valence (OR=1.03, z=0.22, p=0.826) or its interaction with the group factor (OR=0.97, z= –0.14, p=0.885). This suggested that in the “after learning” condition, punishment for advantageous choices did not play a major role in switching to an alternative option in either group and did not have a greater impact on the subsequent “staying or switching” decision in ASD than in NT participants.

Altogether, we observed highly similar performance patterns in both learning conditions between ASD and NT participants. For both NT and ASD participants, we reproduced our previous results obtained for the larger group of NT subjects. This strongly suggested that intellectually high-functioning adults with ASD used typical search strategies associated with performance and task success in the simple two-alternative probabilistic environment. Therefore, atypical phasic pupil response or choice reaction time to exploitative versus explorative decisions in ASD, as we describe in the following sections, was unlikely to be attributed to any performance differences. Indeed, a simple task was chosen specifically to examine whether decision-making challenges in autism arise mainly because of subjective evaluations of one’s decisions that were objectively correct and rational.

### Response time: effects of learning

The linear-mixed effects model of the effects of the Choice Type and Learning on choice response time produced the following statistically significant factors and their interactions: Choice Type (F (1, 4776.7) = 82.4, p < 0.001), Learning (F (1, 4764.5) =6.12, p=0.013), Learning × Choice Type (F (1, 4773.4) = 19.2, p < 0.001). The highly significant Learning × Choice Type interaction suggested that in both groups appearance of the behavioral preference for the advantageous HP over disadvantageous LP choices was paralleled by changes in their relative choice response speed.

As Figure 2 illustrates, in the “no learning” condition, significant modulation of RT by choice type was absent in either group (Tukey HSD for LP vs HP: NT: p = 0.125; ASD: p = 0.079) indicating that both groups made their HP and LP choices with the same speed. In contrast, under the “after learning” condition, the Choice type factor modulated RT in both groups (Tukey HSD for LP vs HP: NT: p <0.001; ASD: p<0.001). In addition, compared with the same types of choices made during a random exploration, i.e., in the “no learning” condition, in the “after learning” condition, response times decreased for HP choices (Tukey HSD: NT = 0.012, ASD = 0.141) and increased for LP choices (Tukey HSD: NT = 0.007, ASD = 0.395).

**Figure 2.**
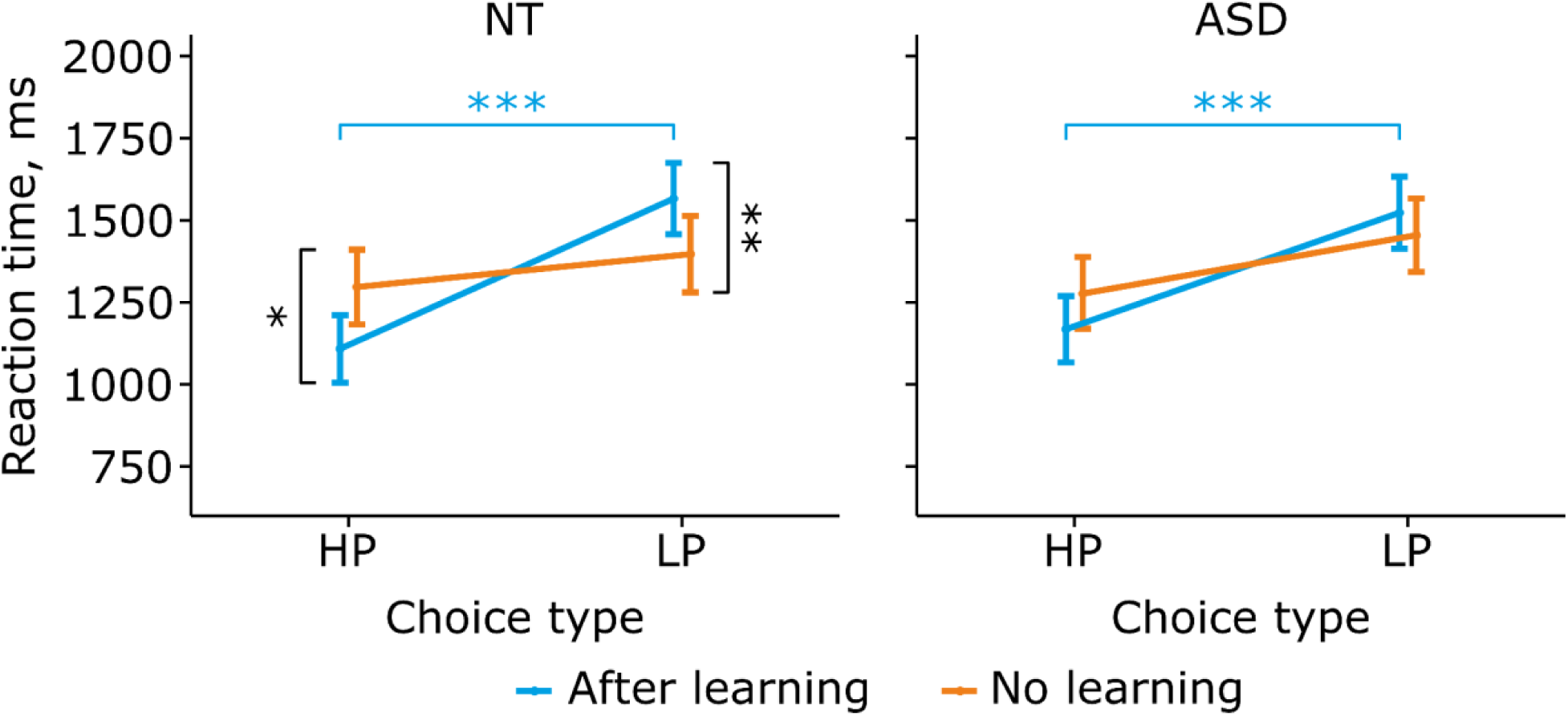
Response time as a function of choice type and learning in NT and ASD groups. RT for HP and LP choice types before (red) and after (blue) the predictive model was learned in NT participants (A) and participants with ASD (B). Points and error bars on graphs represent M ± SEM across single trials in all subjects. Here and hereafter: * p < 0.05; ** p < 0.01; *** p < 0.001

Although the pair-wise comparisons suggested that it was the NT group that mostly contributed to the crossover interaction between Learning and Choice Type, no 3-way interaction with the factor Group was detected (Group*Learning*Choice Type: F (1,3776.5) = 0.92, p=0.337) indicating that the same trend was present in the ASD group as well.

To sum up, in NT and ASD groups alike, the pattern of learning-induced RT changes suggested that outcome predictability facilitated decision-making for objectively advantageous decisions and hindered it for disadvantageous ones by converting them into safe exploitative and risky explorative decisions respectively.

### Pupillary responses

#### Pupillary waveforms in explorative and exploitative choices

First, we aimed to look at the pupil dynamics over the entire trial and at the difference between explorative versus exploitative choices in each group of participants separately (Fig. 3). Overall, pupil dynamics displayed characteristic patterns of dilation and constriction observed in the previous work (Kozunova et al., 2022). The pupil started to dilate after the choice (button press), peaking at more than 1300 ms after the choice (300 ms after a feedback onset), and then constricted with a trough at around 600 ms after visual feedback (Fig. 3). In both NT and ASD participants alike, the pupil size dynamics differed between exploitative and explorative choices. Permutation test with FDR correction revealed that pupil size started to increase on risky explorative choices compared with safe exploitative choices even before the button press and this modulation continued until trial offset. It was detectable as larger pupil dilation in the time interval between choice and feedback and later on as attenuated pupil constriction caused by a light reflex to the visual feedback. The increased dilation and reduced light-induced constriction for more arousing events replicated the pattern found in previous studies and reflected the influence of luminance-independent noradrenergic arousal on the iris dilator muscles (Bradley et al., 2017). In both groups alike, the relative increase in pupil size for risky explorative choices occurred before the feedback was presented suggesting that post-feedback pupil-linked arousal is difficult to distinguish from that caused by a decision to undertake risk. Thus, explorative decisions were more arousing than exploitative ones in ASD and NT participants putatively because they were associated with greater subjective uncertainty in the outcome of a risky choice, which conflicted with the inner predictive model.

**Figure 3.**
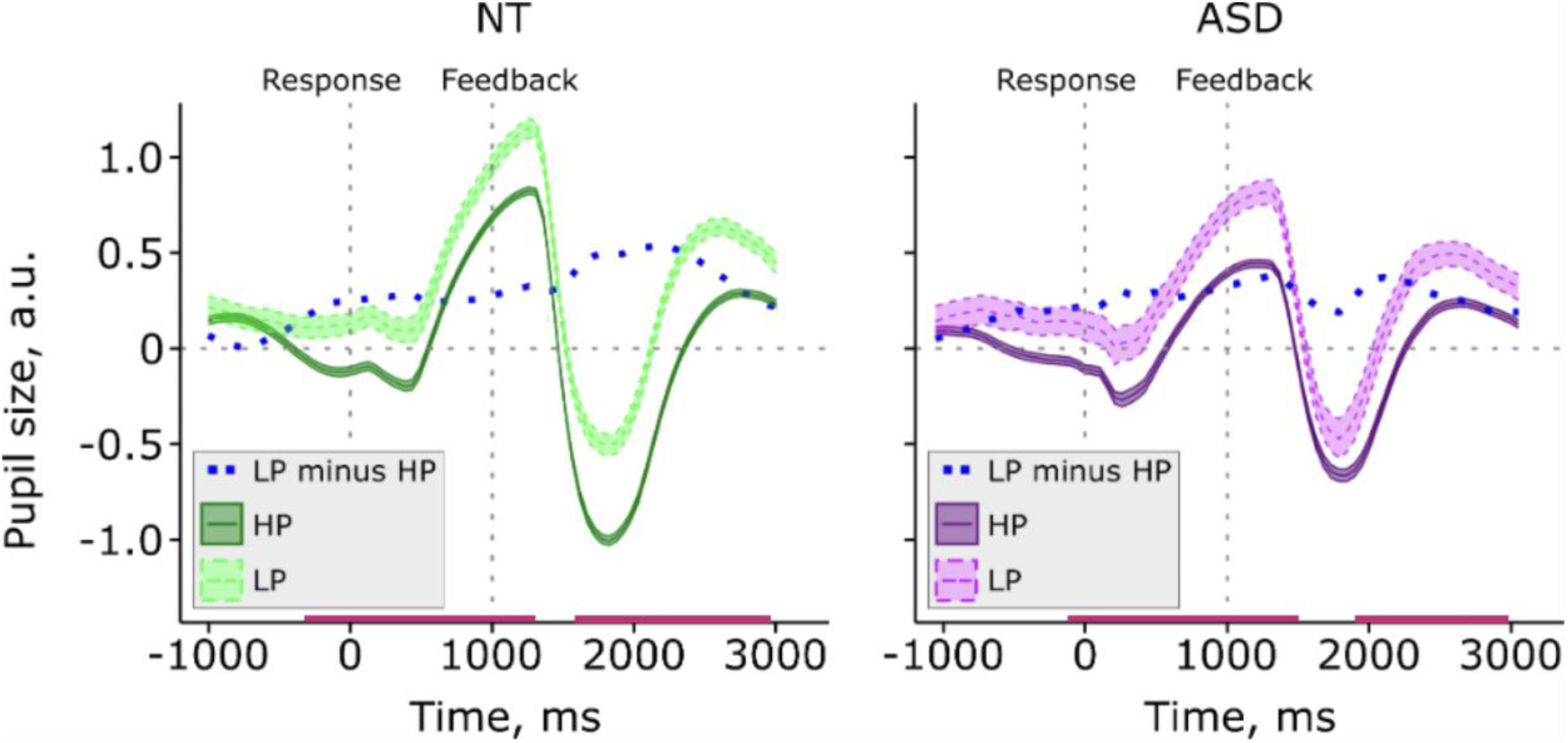
Grand average time courses of pupil size during exploitative HP and explorative LP choices in NT (left panel) and ASD (right panel) participants in the “after learning” condition. Pupil size is within-subject z-scored, and time-locked to button press. Light/dark green and light/dark violet waveforms correspond to LP/HP choices in NT and ASD participants respectively. The dashed curve in each graph represents the time course of the difference in pupil size between the LP choice type and the HP choice. Dashed vertical lines at 0 ms and 1000 ms designate button press and onset of visual feedback respectively. Shaded areas represent M ± SEM across single trials. Horizontal bars at the bottom of each graph mark significant differences between the LP and HP choice types (p < 0.05, FDR corrected).

#### Learning-driven modulations of pupillary responses in ASD and NT

For all further analyses, the pupil size was averaged from 1000 to 2200 ms after the response onset and used as a dependent variable for the LMM analysis with the following fixed effects: Choice Type (HP and LP choices), Previous Feedback (gain and loss), Learning Condition (“no learning” and “after learning”), Group, and their interactions (Table 4 shows the significant effects). Importantly, unlike the RT, pupil response did not show a significant two-way interaction between Learning and Choice Type, yet there was a significant three-way interaction between these two factors and factor Group (F (1, 3788.6) = 10.7, p = 0.001). This suggested that ASD and NT participants had more differences than commonalities in the impact of learning on the pupil response differentiating HP and LP choices.

**Table 4.** Pupillary responses: significant LMM effects.

| Factor | NumDF | DenDF | F | *p* |
| --- | --- | --- | --- | --- |
| Choice Type | 1 | 3799 | 34.5 | <0.001 |
| Learning | 1 | 2622.6 | 5.9 | 0.015 |
| Previous Feedback | 1 | 3783 | 14.7 | <0.001 |
| Choice Type × Previous Feedback | 1 | 3792.7 | 6 | 0.014 |
| Group × Choice type | 1 | 3785.8 | 3.9 | 0.048 |
| Learning × Choice Type × Group | 1 | 3788.6 | 10.7 | <0.001 |
| Learning × Previous Feedback × Group | 1 | 3780.2 | 7.2 | 0.007 |
| Learning × Previous Feedback × Choice type | 1 | 3800.1 | 4.9 | 0.027 |

To analyze the latter interaction effect, we ran the LMM in the “no learning” and “after learning” conditions separately using the model:

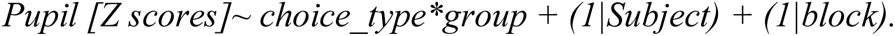

In the “no learning” condition, although participants from both groups did not prefer any alternatives and chose them at the same rate (being involved in random exploration), there still was a significant Choice-Type*Group effect on pupil size (F (1, 513.7) = 8.7, p =0.003).

As Figure 4 illustrates, greater pupil response to LP versus HP choices was significant in ASD participants (ASD, HP versus LP in ‘No learning’, Tukey HSD, p=0.004), while in the NT group, pupil size did not differentiate between HP and LP choices (NT, HP versus LP in ‘No learning’ condition, Tukey HSD, p = 0.427). The absence of a reliable effect of Choice Type on pupil size in NT participants was consistent with our previous finding obtained in a much bigger sample of NT subjects, and thus it cannot be attributed to an underpowered analysis. Thus, the Group*Choice type effect in the “no learning” condition was due to atypically augmented pupil response to objectively disadvantageous LP versus advantageous HP choices in ASD subjects.

**Figure 4.**
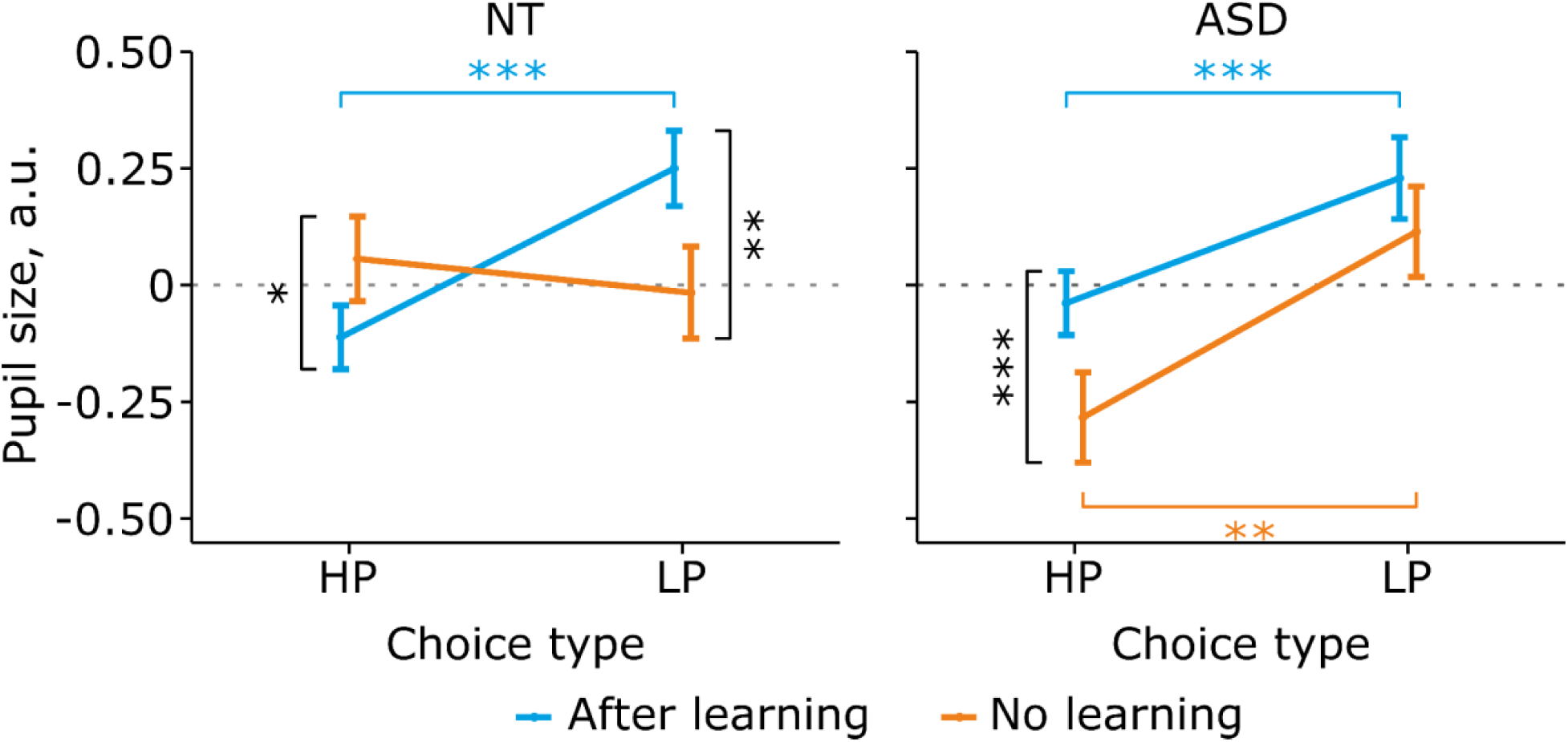
Pupil size as a function of choice type and learning condition. Pupil size (within-subject z-scored) for HP and LP choice types before (orange) and after (blue) the prediction model was learned, in NT participants and ASD participants. Points and error bars on graphs represent M ± SEM across single trials in all subjects. * p < 0.05; ** p < 0.01; *** p < 0.001.

In the “after learning” condition, when participants predominantly (and more quickly) chose an objectively advantageous HP option, i.e., made exploitative choices matching a prediction model, the Choice Type effect became significant in both groups: (NT & ASD samples combined: F(1, 3271.5) = 49.6, p < 0.001). The pupil dilated more during explorative LP choices as compared with exploitative HP ones (HP vs LP, NT: Tukey HSD, p < 0.001; ASD: Tukey HSD, p < 0.001), in line with the hypothesis, which related the enlarged pupil size to heightened subjective uncertainty during predictably risky decisions.

Choice-Type*Group interaction after learning was only trend-level significant (F (1, 3272.1) = 2.8, p = 0.094), but yet deserved consideration because it contributed to between-group differences in the effect of learning on discriminative PDR (three-way interaction effect). After learning, the discriminatory PDR tended to become smaller in ASDs than in NTs instead of being larger as it was before learning (Fig. 4, Suppl. Fig. 1). This suggested that the acquisition of the prediction model shifted group differences in pupillary response in the opposite direction to what was observed for the “no learning” condition of random exploration.

Indeed, direct comparisons of PDR for the same choice type made in the “after learning” vs no learning conditions revealed learning-related changes that differed between the groups for both LP and HP choices (Fig. 4). Regarding LP choice, ASD participants, unlike NT volunteers, did not dilate their pupils more when LP choices became predictably risky (and participants learned that they were associated with low pay-off) than when the same choices represented random explorative strategy (LP choices after vs no learning: NT: Tukey HSD, p=0.004; ASD: Tukey HSD, p = 0.241). The reduced effect of learning for LP choices in ASD participants was likely to be a consequence of atypically high sensitivity of their pupil response to LP choices under the “no learning” condition, rather than being due to decreased response to risky exploratory decisions in the “after learning” condition. However, the PDRs to HP choices in ASD were also associated with atypical learning-related changes. In accord with our previous results (Kozunova et al., 2022), among the NT group, the PDR during HP choices was reduced when participants learned that these choices were associated with a high pay-off (HP after vs no learning: Tukey HSD, p = 0.013). These pupil changes were concordant with the occurrence of a strong preference and shortened decision time for HP choices, thus suggesting that a shift from random exploration to exploitation of a predictive model led to reduced subjective uncertainty in the desirable outcome. Notably, within the ASD group, this difference was in the opposite direction: HP choices drove larger PDRs in “after learning” than in “no learning” conditions (Tukey HSD, p<0.001).

To summarize, the atypical pattern of pupil dilation response in ASD participants was mainly due to two reasons: (i) high sensitivity of pupil response to disadvantageous vs advantageous choices during random exploration (“no learning” condition) that strongly dissociated from the performance measures (choice ratio, RT), which showed no bias in participants’ behavioral preference towards any alternative; (ii) paradoxically increased pupil response to predictably advantageous exploitative HP choices in comparison to the same HP choices made during random exploration (after learning vs no learning contrast). The presence of these atypical features led to the opposite effect of predictive model acquisition on the discriminative pupillary response between exploitative and explorative choices in autistic individuals (negative) compared to neurotypical individuals (positive).

#### The effect of previous feedback in ASD and NT

In line with our previous results for the bigger NT sample, PDR changed during a current choice depending on the valence of the previous feedback (gain or loss). Significant Choice Type*Previous Feedback interaction (F (1, 3792.7) = 6, p=0.013) indicated that the impact of the previous feedback depended on the Choice Type. The interaction was driven by a significant impact of previous gains versus losses on a current-trial PDR, which was selective for the LP choices (Gains vs Losses: LP: Tukey HSD, p < 0.001); HP: Tukey HSD, p = 0.223) (Fig. 5). Yet, this two-way interaction was significantly influenced by the Learning factor (Learning × Previous Feedback × Choice type: F (1, 3800.1) = 4.9; p=0.036). To explain this interaction effect, we ran the LMM on “no-learning” and “after-learning” data subsets in the combined sample of NT and ASD participants. Choice Type × Previous Feedback interaction was significant within the after-learning condition only (no learning: F (1, 514.6) = 0.01, p=0.920; after-learning: F (1, 3262.8) = 18.4, p <0.001). When a prediction model was learned, in both groups alike, the pupil size for LP choices was greater after previous gains compared with losses (NT: ‘Losses’ vs ‘Gains’ for LP, Tukey HSD, p <0.001; ASD: ‘Losses’ vs ‘Gains’ for LP, Tukey HSD, p < 0.001). At the same time, for HP choices the difference was either absent (NT: ‘Losses’ vs ‘Gains’ for HP, Tukey HSD, p = 0.174); or greatly reduced (ASD: ‘Losses’ vs ‘Gains’ for HP, Tukey HSD, p = 0.037). This suggested that ASD and NT participants alike, dilated the pupil more when shifted their action plan toward exploration (LP-choice), but especially so for those explorative choices, which they made after being rewarded for the immediately preceding exploitative action (i.e., HP-choice). In other words, among ASD and NT participants, the maximal PDR was observed for those explorative choices that conflicted not only with the accumulated trial-and-error experience but also with the most recent trial outcome.

**Figure 5.**
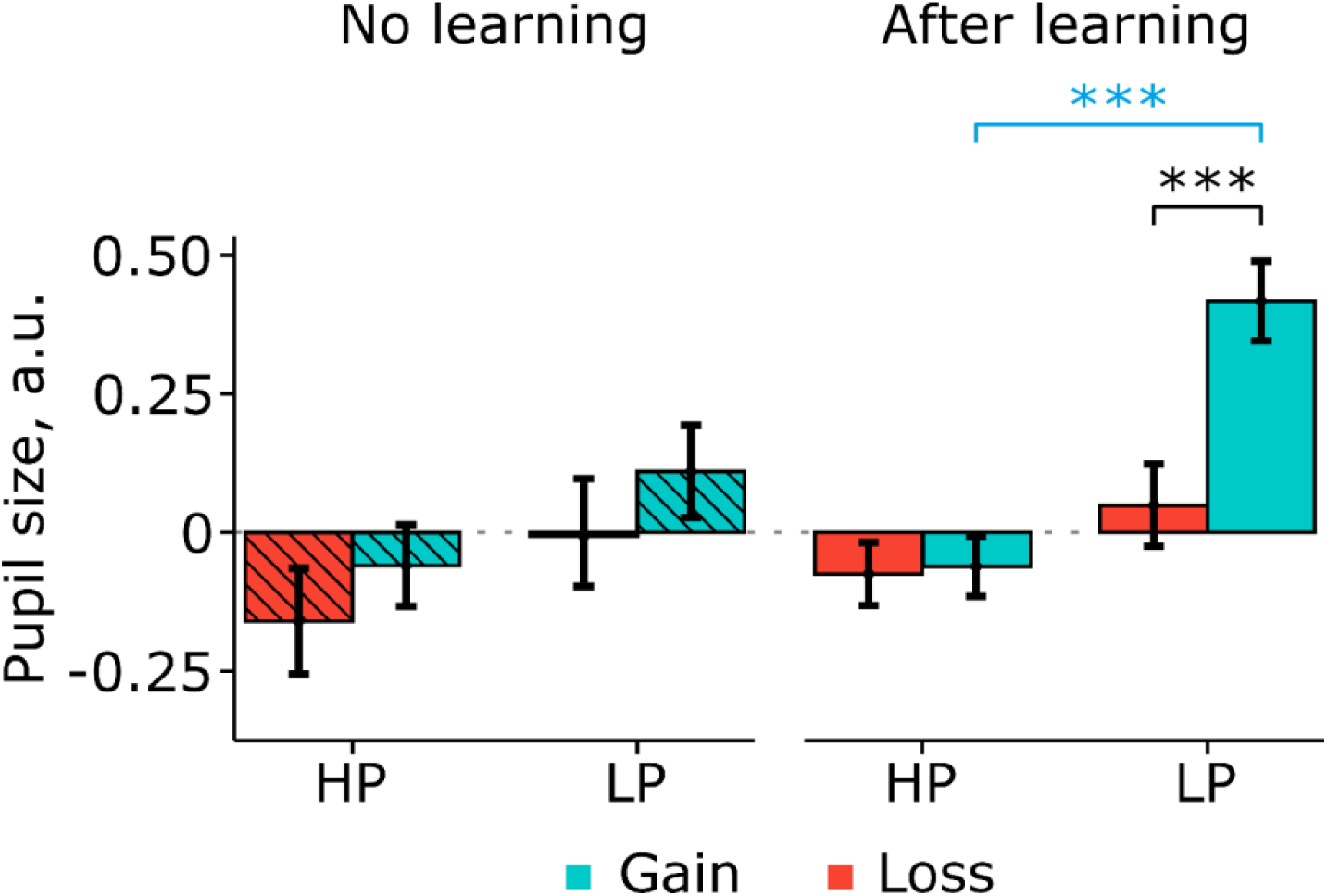
Effect of Previous Feedback on pupil dilation response under “no learning” and “after learning” conditions in the combined NT & ASD sample. Turquoise: previous gains; Salmon: previous losses, all the other designations as in the previous Figures.

A significant interaction effect of Group*Learning*Previous Feedback was mainly explained by a greater learning-induced difference in PDR resulting from previous positive versus negative feedback in ASD than NT participants regardless of choice type (Suppl. Fig. 2).

#### Effect of the Current Feedback

In addition, we considered the putative impact of the positive versus the negative outcome of a current choice on PDR. A respective LMM model with fixed factors Learning, Choice Type, Group, and Current Feedback valence revealed neither the main effect nor any significant interactions with the factor of Current Feedback. Thus, as far as we can tell from our data, the differential PDRs we observed within the post-feedback intervals did not reveal any visible relation to the outcomes of current decisions; seemingly, at least a substantial portion of the effect during the post-feedback interval was just a continuation of differential post-response effects, which were slowly developing in time and extended well into the post-feedback time interval.

#### Intolerance of uncertainty and its correlations with pupillary responses

ASD compared with NT participants demonstrated significantly higher levels of self-reported Intolerance of Uncertainty (Mann–Whitney U =307, n1=23, n2 = 17, p=0.002, two-tailed; see Fig. 6). This finding is in line with previous studies (for meta-analysis see (Jenkinson et al., 2020)) showing increased Intolerance of uncertainty in ASD.

**Figure 6.**
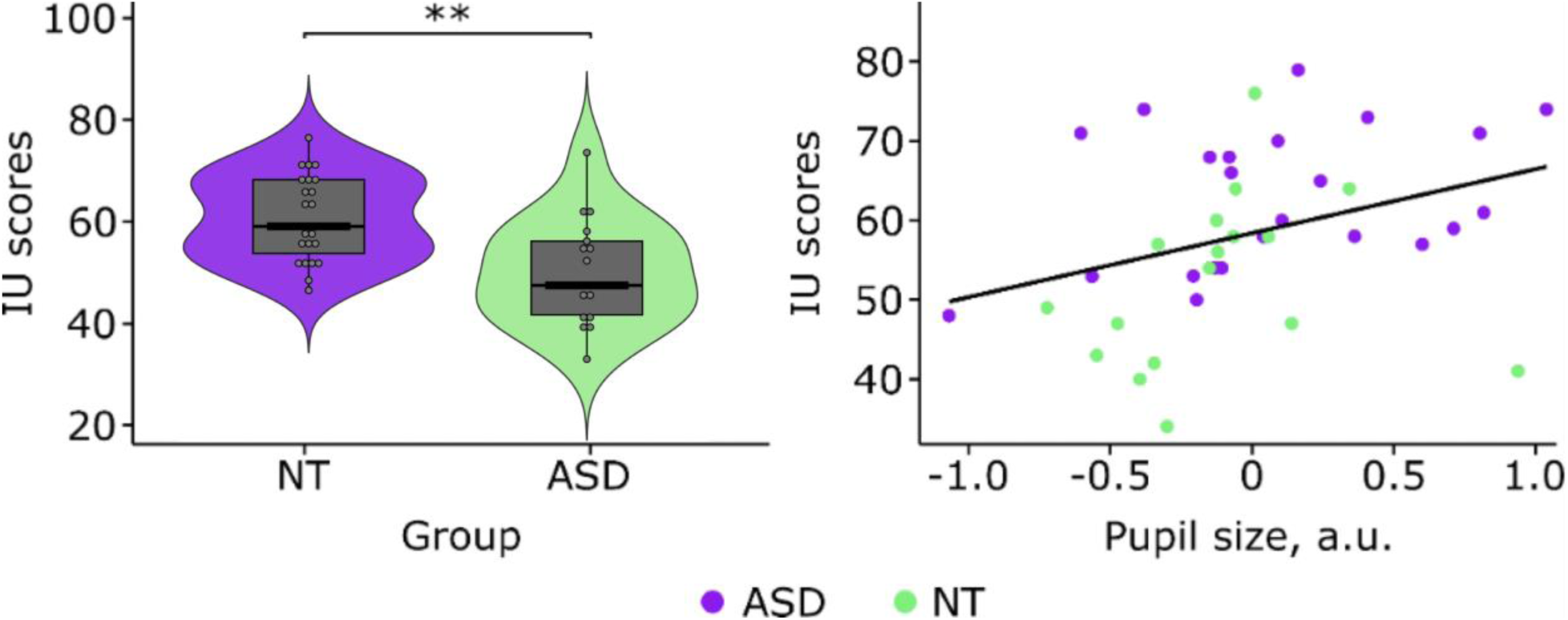
Intolerance of uncertainty in NT and ASD participants (left panel) and its correlation with PDR induced by exploitative choices (right panel). The line on the scatterplot represents linear regression for the combined sample.

Spearman correlations between IU scores and pupil size in LP and HP choices under the “after learning” condition in ASD and NT groups are presented in Table 5 and illustrated in Fig. 6.

**Table 5.**
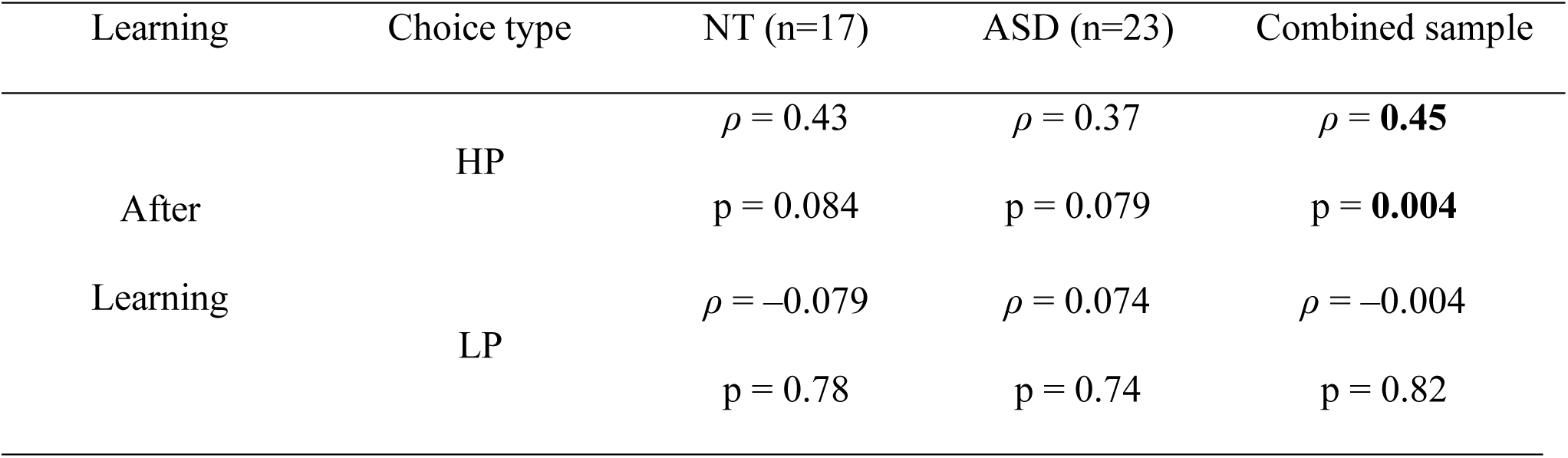
Spearman’s correlations between choice-related PDR and Intolerance of Uncertainty. Spearman correlation coefficients (uncorrected) are shown, and correlations that survived after the Bonferroni correction are highlighted in bold. The reduced number of NT participants is due to missing data on the IU questionnaire.

The results showed one significant correlation, which survived after Bonferroni correction for multiple comparisons. Greater pupil dilation during safe exploitative choices (i.e., HP) characterized participants with higher IU scores in the pooled sample of 40 subjects (ρ (38) = 0.45, Bonferroni corrected p = 0.028)). The same trend was observed in ASD and NT groups separately, although it did not reach an accepted significance level in either group.

Although we did not have a hypothesis for the parallel relation with Intolerance of Uncertainty involving PDRs to HP and LP choices in the “no learning” condition, we looked for their presence and found no significant correlations at the uncorrected p-level of 0.05 (Suppl. Table 1)

Thus, higher intolerance of uncertainty in our participants was particularly associated with greater pupil response to the exploitative choices, which had the lowest estimated probability of negative outcomes and were supposedly considered the least stressful events among the four types of choices the participants made throughout the experiment.

## Discussion

In ASD and NT subjects, we analyzed behavioral and pupillometric concomitants of decision-making under uncertainty when participants learned by trial and error to discriminate between objectively advantageous options (HP, high reward probability) and disadvantageous options (LP, low reward probability) in a simple two-alternative reward learning task with a stable probabilistic reward structure (70 and 30% reward probabilities for HP and LP options, respectively). The findings show that in participants with ASD, their seemingly normal ability to learn to predict the outcome of choosing the best option among alternatives contrasts sharply with the atypical effect of prediction learning on pupil-related arousal. Moreover, it is in preferred HP choices with predictably better outcomes that greater pupil-related arousal directly correlates with self-reported intolerance of uncertainty in daily life. The dissociation between rational decisions and their autonomic concomitants in a stable probabilistic environment is a novel finding that provides additional insight into why individuals with autism have difficulties dealing with uncertainty.

Consistent with the previous reports (Hwang et al., 2020; Vasa et al., 2018), our participants with ASD, whose performance IQ did not differ from the neurotypical sample, had higher scores on the Intolerance of Uncertainty Scale than the NT participants (Fig. 6). However, their predictive abilities in a stable probabilistic environment were not disturbed. Subjects with ASD did not require more trials to achieve the performance criteria and were no worse than NT subjects in adjusting their task strategy and preferentially selecting advantageous stimuli after choice-reward contingency was learned and a choice value could be predicted (Table 3). Also, similarly to NT subjects, they chose the advantageous stimuli faster than disadvantageous ones after – but not before – reaching the learning criteria, indicating that an exploitative decision is easier to make than its explorative alternative, which contradicts the prediction of its low value (Fig. 2). The finding that uncertain or conflictual decisions are slower than decisions for which more information is available is common in the decision-making literature (Zenon, 2019). Thus, participants with ASD were as efficient as NT participants in internalizing prior information (inner predictive model) to make predictions about an upcoming choice outcome and use these predictions to guide their decisions. Our behavioral results confirm that when choice and outcome contingency are stable, individuals with ASD adapt their behavior similarly compared with NT subjects.

In sharp contrast to typical reward-learning behavior, pupil-linked arousal accompanying decision-making exhibited more differences than commonalities between ASD and NT individuals.

Unlike NT, subjects with ASD exhibited a discriminative pupil response to disadvantageous choices when they chose alternative options with equal probability and equal speed, i.e., in the “no-learning” condition (Fig. 3 and Fig. 4). The augmented pupil response to disadvantageous compared with advantageous choices during random exploration was a distinguishing characteristic of the ASD as it was absent in the NT sample. There are two putative explanations of this finding. The first one – the inner prediction model was formed faster in ASD than in the NT, but subjects with ASD needed more external evidence confirming its validity before they transformed it into an overt behavioral preference for advantageous options. The available data speaks against this possibility. Compared with the NT, the ASD group did not need more trials to reach the learning criteria in their task performance, nor did they display a speeded response during HP versus LP choices before the learning criteria were reached (Fig. 2). The presence of a prediction model in the “no learning” condition in ASD subjects is also at odds with the lack of pupil sensitivity to immediate external evidence of predictive model validity. In both the ASD and NT groups, only under the “after learning” condition, the positive versus negative outcome of HP choice significantly increased pupil dilation for a following exploratory LP choice (Fig. 5). This indicates that a match between a model prediction (when a model exists) and the actual consequences of the preceding HP choice further hampers switching to an exploratory/risky choice making exploratory decisions even more arousing. Therefore, the lack of this effect in the ASD group under the “no learning” condition adds to other pieces of evidence of an absent prediction model that can be used to guide actions.

Yet an alternative possibility is that a relatively more dilated pupil in the LP versus HP choices in the “no learning” condition reflects a negative emotional state associated with such decisions in many previous trials that may dissociate from explicit conscious knowledge about risk. Pupil dilation in response to stimuli that were linked to threat in the life-long experience has been reported as an indicator of increased emotional arousal (Bradley et al., 2008; Henderson et al., 2018), and its neural origin is distinct from that of pupil response induced by expectancy, uncertainty, surprise, or other cognitive processes (Critchley et al., 2001). In ASD, previous evidence in the literature on atypically enhanced autonomic arousal caused by probabilistic negative reinforcement comes from the field of fear conditioning of skin conductance response (SCR). In the probabilistic paradigm, adults with ASD being compared with typical individuals, display augmented SCR responses to stimuli associated with threat during the conditioning procedure (Espinosa et al., 2020). Somatic marker theory (Damasio, 2004) proposes that in certain contextual situations, memories of body states associated with previous feeling states automatically trigger both autonomic responses and positive or negative emotions (brain-body loops). Typically, the body feeling contributes to future planning and decision-making (Bechara et al., 2005; Mayer, 2011). However, emotional and interoceptive signals are known to have a lesser impact on the decision-making process in ASD adults without cognitive impairment (Shah et al., 2016). A dissociation between exaggerated negative internal sensations associated with disadvantageous decisions (measured through hypersensitive pupil response) in combination with reduced awareness of their feelings in ASD, may explain why their “premorbid” autonomic sensitivity did not improve prediction abilities and was not transformed into a prediction model. This explanation is also consistent with the idea that, instead of using interoceptive or emotional information during their decision-making, individuals with ASD use exclusively a rule-based strategy (Brosnan et al., 2016; Shah et al., 2016). Apparently, similar post-decisional pupil dilations can index different processes depending on the context in which a decision is made.

Indeed, when learning occurs and participants begin to base their decisions on predicted value of respective choices, exploratory/risky decisions in both ASD and NT individuals are more arousing than exploitative/safe decisions (Fig. 3 and 4). This finding replicates the previous results obtained in the bigger sample of neurotypical individuals (Kozunova et al., 2022) and agrees with the notion that upon a prediction model acquisition, pupil size changes are mainly triggered by cognitive processes associated with using/updating the existing model (Zenon, 2019) and are likely mediated through feedback from the prefrontal cortex (Dayan, 2012; Joshi et al., 2016). The increase in pupil size occurs before the external feedback, around the time of decision completion (indicated by button press), and persists a long time after feedback is presented. This means that larger pupil size is not only indicative of the arousing properties of salient feedback, which is scaled with outcome uncertainty (Kreis et al., 2023; Lawson et al., 2017; Pajkossy et al., 2023), but also reflects increased anticipatory subjective uncertainty about the favorable outcome of an exploratory choice that contradicts the model-based prediction of its low value.

Although in the “after learning” condition, there was no more significant difference between groups in PDR, there was a trend towards *less* discriminative pupil response to risky LP explorative versus safe HP exploitative decisions in the ASD group (Fig. 4, Suppl. Fig.1) and it was exactly the opposite of that seen for PDR differentiating LP and HP decisions during random trial-and-error search (in “no learning” condition). In this respect, our finding reminds literature evidence for less discriminative pupillary responses in people with ASD in a variety of conditions related to prediction learning: habituation to standard stimuli in a classic oddball task (Zhao et al., 2022), decision-making in an unstable environment either during an associative perceptual learning task (Lawson et al., 2017) or a probabilistic reversal learning task (Kreis et al., 2023). In all these tasks, the more unexpected (the less predictable in terms of the prior experience) events were accompanied by greater pupil dilation in NTs, and to a lesser extent this was manifested in ASDs. Blunted differential pupillary responses to feedback cues in volatile versus stable environments in autism have been interpreted as an indicator of atypically reduced surprise, or, more specifically, as atypically mild surprise at both predictable and unpredictable outcomes (Lawson et al., 2017). Under our experimental settings, pupil size modulations by outcome predictability appear to reflect differences between ASD and NT individuals at both stages of the decision evaluation process: the anticipation of the impending outcome and the actual response to the outcome – providing that differential modulation of pupil size is present both before and after receiving the feedback signal. From this perspective, the atypical pattern of pupil-linked arousal in ASD subjects is better explained in terms of reduced differences in subjective uncertainty about the future success of their exploratory versus exploitative actions.

Differences between ASD and NT individuals in the modulation of pupil-related arousal by outcome predictability were particularly pronounced when comparing formally identical decisions, e.g., such as those that were objectively advantageous, but made when participants either knew or did not know what to expect from their choice (i.e., during “after learning” and “no learning” conditions respectively).

Surprisingly, the learning-induced changes in pupil-linked arousal were concordant with behavioral measures (RT and choice ratio) in NT but not in ASD subjects (Fig. 3, 4). In NT participants, high-risk explorative LP decisions as compared with the same disadvantageous LP decisions during random exploration were more difficult, i.e., occurred more rarely, took a longer time to make, and concomitantly were more arousing/stressful, i.e. induced greater pupil-linked arousal. Although subjects with ASD exhibited the same behavioral changes, the effect of predictive learning on their LP-related pupillary responses was nonsignificant, probably because, as we discussed earlier, pupil size was already high in the absence of the prediction model during random exploration. On the other hand, the opposite direction of typical learning-induced changes in their pupillary responses to advantageous HP choices most likely stems from a different source. Given that the occurrence of the prediction model not only creates a strong propensity to make exploitative HP decisions more frequently (Table 2) but also speeds them up (Fig. 2), both ASD and NT participants likely perceived such decisions as relatively easy to make. Although in the NT group, the exploitative HP decisions were also less arousing than HP decisions made during random exploration, in subjects with ASD these easy decisions were accompanied by relatively increased pupil-related arousal.

Taken together, these results suggest that an atypical effect of predictive learning on pupil-related arousal in subjects with ASD is predominantly associated with choices for which the expected outcome is good but still uncertain because of the irreducible risk of a probabilistic choice-outcome association. We hypothesized that adult subjects with ASD may be less confident in the inner predictive model than NTs and thus they overestimate the likelihood of a negative outcome for exploitative model-congruent choices, even though they had ample opportunities to learn that this rarely happens. Exaggerated subjective uncertainty does not interfere with their adaptive behavior in our simple task, but causes atypically elevated autonomic arousal when they make choices with predictably high pay-off. On the other hand, both people with NT and people with ASD experience high subjective uncertainty (and high pupillary arousal) when faced with novel situations for which they have no relevant experience or insufficient information about the future outcome of their choices. As a result, the excessive pupillary arousal caused by negative attitudes toward uncertainty in individuals with ASD may be particularly prominent at low levels of uncertainty, attenuated for advantageous choices by prediction model acquisition.

This assumption is supported by direct correlations between inter-individual variations in pupil-related arousal and measures of uncertainty intolerance in our sample (Fig. 6, Table 5). Consistent with the previous reports (Jenkinson et al., 2020), our participants with ASD, had higher scores on the Intolerance of Uncertainty Scale than the NT participants (Fig. 6). When we assessed the PDR at the individual level, its correlation with the uncertainty intolerance trait in the combined ASD & NT sample was related specifically to safe exploitative choices, being absent for the exploratory choice type with high levels of subjective uncertainty (i.e., directed exploration) (Table 5). It is possible that only some patients with ASD, such as those with high levels of uncertainty intolerance, exhibit exaggerated pupillary responses to exploitative/safe decisions. Moreover, the above association is not specific to the ASD diagnosis but becomes more pronounced when a sample includes individuals whose uncertainty intolerance is atypically increased and combined with other psychological features of autism.

Our finding of an aberrant autonomic response to events with a high probability of positive outcomes in individuals with a high intolerance of uncertainty is not unique. Elevated IU is considered a transdiagnostic sign and is not associated with any specific psychiatric diagnosis (McEvoy et al., 2019). It is thought to contribute to anxiety and underlies exaggerated negative affect and physiological arousal in many mental disorders including ASD. Several studies using different methods, paradigms, and patient populations have reported results that were generally reminiscent of our findings. In line with our pupil data in ASD in a simple probabilistic learning task, verbal reports of people with a general anxiety disorder in the same task evidenced that they underestimate the probability of desirable outcomes especially when its true likelihood is high (LaFreniere & Newman, 2019). Moreover, these unjustified worries extended to real life, where such worries did not come true in a majority of carefully monitored outcomes (LaFreniere & Newman, 2020). Regarding autonomic measures, patients with panic disorder who had high IU demonstrated indiscriminate autonomic responding (startle blink) between the cues predicting safety and the cues indicating a following threat of an electric shock (Gorka et al., 2014). Mirroring our pupil results, the results of Gorka and colleagues (Gorka et al., 2014) showed that blunted differential startle response in individuals with high IU was due to potentiated startle to safety cues rather than a decreased response to threat cues.

## Conclusion

Although in our simple two-alternative probabilistic task subjects with ASD, similarly to neurotypical control subjects, learned to reject the events with low value and to prefer the events with high value, there was still atypically heightened pupil-related arousal accompanying their preferred choices, which exploit their prior knowledge of choice outcomes. This aberrant autonomic response to a decision with the best possible but still uncertain outcome was directly related to a greater degree of self-reported intolerance of uncertainty in their daily lives. The atypical pupil-linked arousal cannot be explained by stress triggered by volatile environments and/or inflated estimation of environmental volatility, because the reward structure was stable throughout the experimental session. We interpret the results in terms of heightened subjective uncertainty surrounding current beliefs regarding the best possible choice, i.e., uncertainty in the inner predictive model in ASD. Thus, taken together, our results suggest that difficulties in dealing with environmental uncertainty in ASD subjects are not solely a cognitive problem, but also one of non-confidence in their otherwise unharmed ability to predict the high value of certain options in probabilistic settings.

## Acknowledgments and Funding

We wish to thank V.A. Medvedev for valuable contribution into data analysis.

## Data availability

Data are openly available at 10.6084/m9.figshare.25586199. The experiment was not preregistered.

## Notes

https://figshare.com/s/b5796bf23e83437ebbfe

### Competing Interest Statement

The authors have declared no competing interest.

