## Supplementary for "Atypical Pupil-Linked Arousal Induced by Low-Risk Probabilistic Choices, and Intolerance of Uncertainty in Adults with ASD"

### Supplementary materials

**Supplementary Table 1**

*Spearman's correlations between choice-related PDR and Intolerance of Uncertainty in the 'No learning' condition*

| Learning | Choice type | NT (n=17) | ASD (n=23) | Combined sample |
|---|---|---|---|---|
| No Learning | HP | $\rho$ = –0.02<br>p = 0.947 | $\rho$ = –0.19<br>p = 0.502 | $\rho$ = –0.20<br>p = 0.333 |
| | LP | $\rho$ = –0.16<br>p = 0.681 | $\rho$ = –0.03<br>p = 0.920 | $\rho$ = –0.05<br>p = 0.830 |

**Supplementary Figure 1**

*Between-group comparison of pupil size as a function of choice type and learning condition.*

Points and error bars on graphs represent M ± SEM across single trials in all subjects.

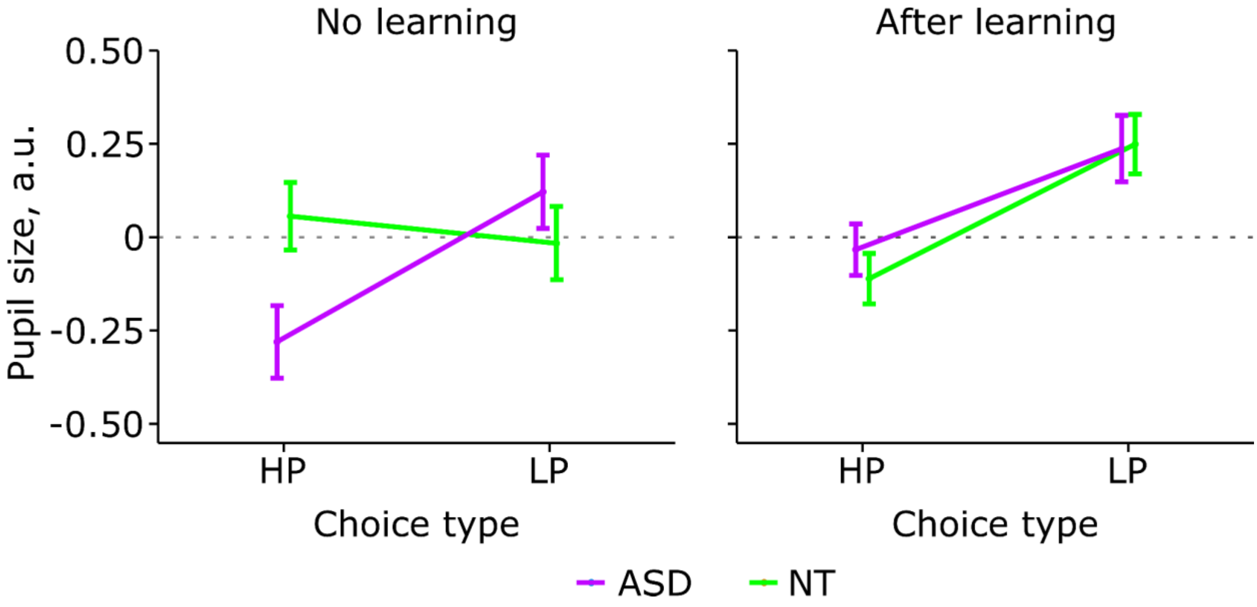

**Supplementary Figure 2**

*Impact of learning and previous feedback on the current-trial PDR in NT participants; and ASD participants*. HP and LP choices are pooled. Turquoise: previous gains; Salmon: previous losses. All the other designations are as in previous Figures.

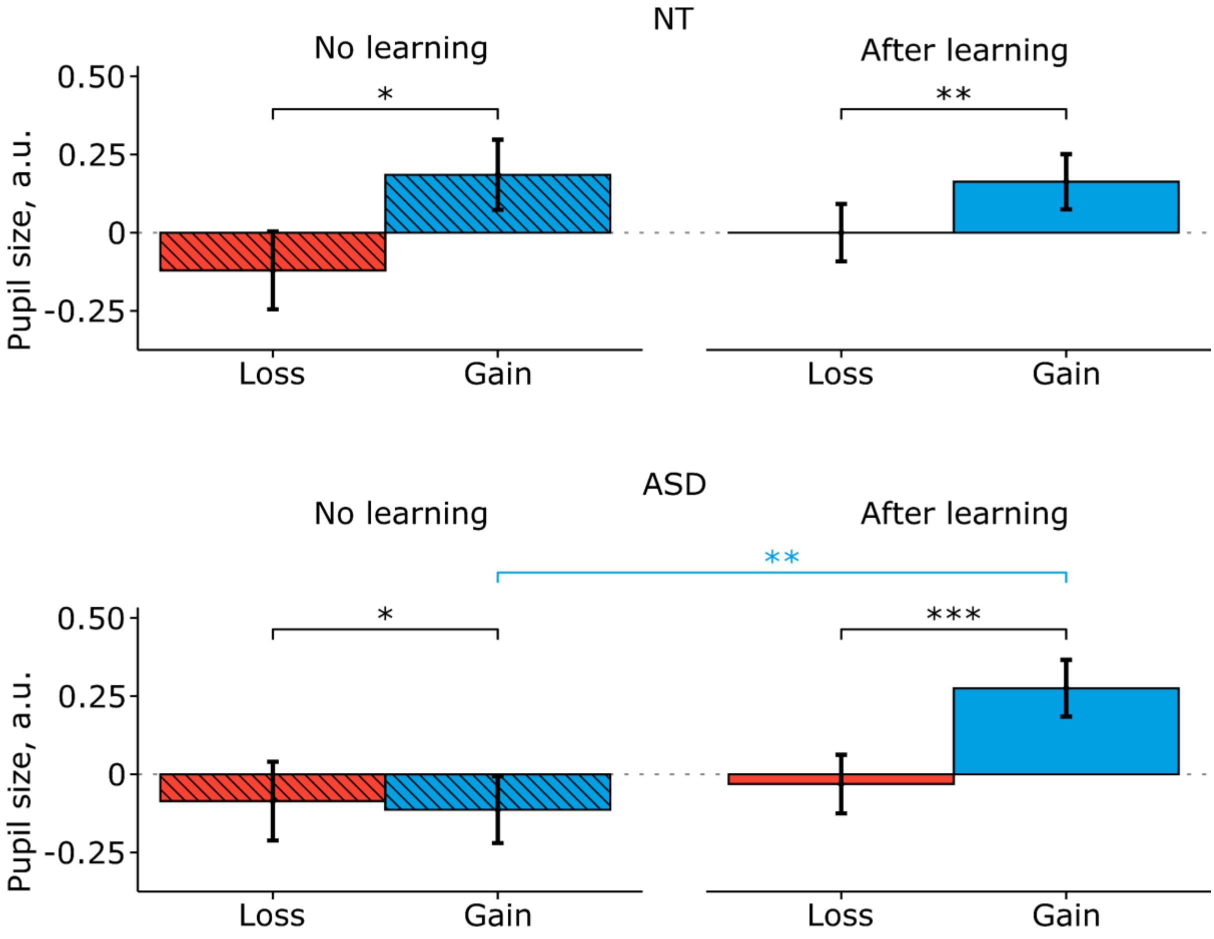
